# Optimization and model averaging of histogram-based place cell firing rate maps using the point process framework

**DOI:** 10.1101/2024.07.05.602233

**Authors:** Murat Okatan

## Abstract

**Background:** The firing rate of hippocampal place cells depends on the spatial position of the organism in an environment. This position dependence is often quantified by constructing spike-in-location and time-in-location histograms, the ratio of which yields a firing rate map.

**New Method:** The purpose of this study is to present a new method for optimizing the spatial resolution of histogram-based firing rate maps.

**Results:** It is pointed out that histogram-based firing rate maps are conditional intensity functions of inhomogeneous Poisson process models of neural spike trains, and, as such, they can be optimized through model selection within the point process framework.

**Results:** The point process framework is used here for optimizing the size and the aspect ratio of the histogram bins using the Akaike Information Criterion (AIC). It is also used for model averaging using Akaike weights, when maps of various bin sizes provide comparable fits. Application of the method is illustrated on data from real rat hippocampal place cells.

**Comparison with existing methods:** Existing methods do not optimize the number of bins used in each dimension of the firing rate map.

**Conclusion:** The proposed approach allows for the construction of the AIC-best histogram-based firing rate map for each individual place cell.

## 1. Introduction

Hippocampal place cells were discovered by O’Keefe and Dostrovsky while they were recording the spiking activity of neurons using microelectrodes implanted into the dorsal hippocampus of rats (O’Keefe & Dostrovsky, 1971). The cells seemed to respond primarily to the spatial location of the animal in the experimental environment. Subsequent studies examined this putative position dependence with increasing systematicity (Muller et al., 1987; Muller & Kubie, 1989; O’Keefe, 1976; O’Keefe & Conway, 1978).

One of the early methods of quantifying the position dependent firing of place cells was a histogram-based firing rate map (Muller et al., 1987; O’Keefe, 1976). In this method the spatial coordinates of a rat’s head are tracked using a video camera, and the spiking of individual neurons is recorded simultaneously with the position data. In this way, the location at which a spike is observed can be determined. The experimental environment is divided into adjacent non-overlapping and contiguous spatial bins. The number of spikes fired by a cell is determined for each bin (spike-in-location map). The total amount of time spent in each bin is also determined (time-in-location map). The firing rate map of the cell is then determined as the ratio of the spike-in-location map to the time-in-location map (Muller et al., 1987).

In early studies, the size and the aspect ratio of the bins were determined by the area viewed by each pixel of the camera and by how many pixels were used per bin, which resulted in bin sizes such as 3.5 (X) by 2.8 (Y) cm (Muller et al., 1987; Muller & Kubie, 1989). In more recent studies, square bins of edge length 5 cm (Bures et al., 1997; Solstad et al., 2014; Tanila et al., 2018), 4 cm (Schoenenberger et al., 2016), 3 cm (Fenton & Muller, 1996, 1998; Harvey et al., 2020; Hayman et al., 2015; Ismakov et al., 2017), 2.3-2.7 cm (Jung et al., 1994; Karnam et al., 2009; Muller, 1996; Muller et al., 1994; Zhang et al., 2014; Zhou et al., 2007), 2 cm (Sakkaki et al., 2020; Skaggs & McNaughton, 1998; Wikenheiser et al., 2021), 0.9-1 cm (Ásgeirsdóttir et al., 2020; Hazama & Tamura, 2019), 0.5 cm (Grieves, 2023) have been used. In these studies, a fixed bin size is used for all cells. Yet, when the firing rate assigned to a bin is viewed as a model parameter, using the same number of bins for all cells may overfit the data of some cells, while it may underfit the data of other cells. Thus, it is desirable to optimize the number of bins used in a firing rate map, separately for each cell, and in a fully data-driven manner, which would also optimize the size and the aspect ratio of the bins.

While the optimization of the firing rate estimates in each bin of a firing rate map has been the subject of some earlier studies (Jung et al., 1994; Skaggs et al., 1996; Yartsev & Ulanovsky, 2013), none of those studies optimized the number of bins contained in the firing rate map, nor did they use the point process framework. Therefore, here, this framework is used for optimizing the size and the aspect ratio of the firing rate map bins, and the optimal firing rate estimates per bin are obtained directly as a result of this optimization. In the method presented here, the size and the aspect ratio of the bins are model parameters that are optimized using the Akaike Information Criterion to obtain the best fit to the available data. The proposed method is also used for computing model-averaged firing rate maps using Akaike weights when two or more maps provide comparable fits to the data. Application of the method is illustrated on data from real rat hippocampal place cells. Advantages of using the proposed method in place of existing firing rate construction methods are discussed.

## 2. Materials and methods

### 2.1 Data

The present analysis uses the place cell data published by Cajigas (2017). These data were collected from two Long-Evans rats that were freely foraging for randomly delivered food pellets in an open circular environment of 70 cm diameter with 30 cm high walls and a fixed visual cue (Barbieri et al., 2005; Brown et al., 1998; Cajigas et al., 2012). The rats were well familiarized with the environment prior to data collection, which implies that place fields were formed and stabilized to a large extent. Simultaneous activity of 49 (37) place cells was recorded for 24 (23) min using a microelectrode array that was implanted into the CA1 region of the hippocampus of rat 1 (rat 2). The sampling rate was 31.25 kHz per electrode. The position of the rat’s head was measured at 30 Hz by a camera tracking the location of 2 infrared diodes mounted on the head stage.

## 3. Theory and calculation

This section explains how the histogram-based firing rate map analysis of place cell spiking is formulated and shows that the firing rate map is equivalent to the conditional intensity function of an inhomogeneous Poisson process model of neural spiking. Optimization of this model is explained.

### 3.1 The Firing Rate Map

To quantify the position-dependence of place cell spiking, Muller et al. (1987) conducted a by now classical experiment in which rats freely foraged for randomly delivered food pellets in a cylindrical environment with 76 cm diameter and 51 cm high walls. During the experiment, they recorded the position of the rat’s head simultaneously with hippocampal neural activity. The animal’s head position was tracked using a TV camera. By isolating the spike trains of individual hippocampal neurons, they were able to identify the locations at which each individual cell fired spikes. Although similar video/computer methods for locating the animal were exployed in previous studies (McNaughton et al., 1983; O’Keefe, 1983), 2-D histogram-based construction of the firing rate maps was explained in detail first in (Muller et al., 1987).

Let 0 ≤ *t*_1_ < *t*_2_ < … < *t*_*N*(*T*)_≤ *T* denote the spike times of a place cell, where *N*(*T*) is the total number of spikes recorded from that cell throughout the experiment, and *T* denotes the duration of the recording. Let 0 ≤ *τ*_1_ < *τ*_2_ < … < *τ*_*K*_ ≤ *T* denote the position sampling times and (*x*(*τ*_*k*_), *y*(*τ*_*k*_)) the location of the animal’s head at the time of the *k*^th^ position sample.

Muller et al. divided the image captured by the camera into a 64×64 array of non-overlapping pixels (bins) to generate spikes-in-location and time-in-location arrays (Muller et al., 1987). The circular environment corresponded to a 23 (X) by 28 (Y) array of bins. The value of a bin in the spikes-in-location array indicated the number of spikes fired by a neuron in that bin, whereas the value of a bin in the time-in-location array indicated the total amount of time the animal spent in it. The firing rate map was obtained by dividing the spikes-in-location array by the time-in-location array. Bins that were not visited by the animal were ignored.

#### 3.1.1 Expressing the Firing Rate Map as a Conditional Intensity Function

This section shows that the firing rate map of Muller et al. (1987) models a place cell’s spike train as an inhomogeneous Poisson process with the conditional intensity function given in Eq. 1

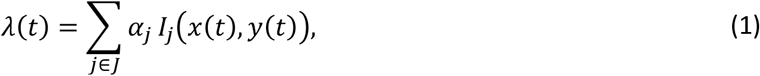

where (*x*(*t*), *y*(*t*)) is the instantaneous position of the animal, *I*_*j*_(*x*(*t*), *y*(*t*)) is 1 if the position is inside the *j*^th^ bin and 0 otherwise, and *α*_*j*_ is the rate of spiking inside the *j*^th^ bin. In this way, the spiking of a place cell is modeled as a homogeneous Poisson process within each bin, but the rate is allowed to change from one bin to another, making the overall model the conditional intensity function of an inhomogeneous Poisson process. *J* is the set of all bins that are visited by the animal during the experiment. Note that the functions *I*_*j*_(*x*(*t*), *y*(*t*)) form an orthonormal basis since the bins are non-overlapping.

Let the firing rate map be an *m*-by-*n* array of non-overlapping adjacent and contiguous bins, where the bin at row *r* and column *c* is indicated by the pair (*r,c*). For 0 ≤ *t* ≤ *T*, let 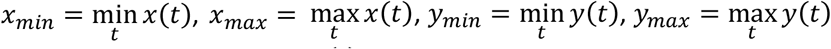 denote the minimum and maximum values that *x*(*t*) and *y*(*t*) processes attain during the experiment, respectively.

Let the row number corresponding to *x*(*t*) be computed as

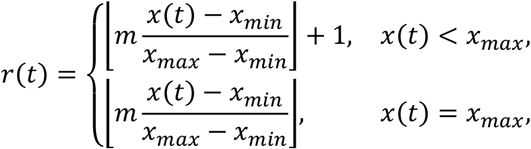

and the column number corresponding to *y*(*t*) be computed as

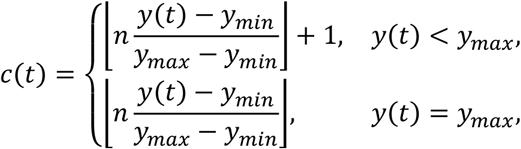

where ⌊·⌋ denotes the floor function, which returns the largest integer that is smaller than or equal to its argument.

Using these row and column number definitions, *I*_*j*_(*x*(*t*), *y*(*t*)) = 1 if *j* = *r*(*t*) + (*c*(*t*) − 1)*m*, and *I*_*j*_(*x*(*t*), *y*(*t*)) = 0 otherwise.

The log-likelihood function of the parameter vector *α* is (Daley & Vere-Jones, 2003, p. 233)

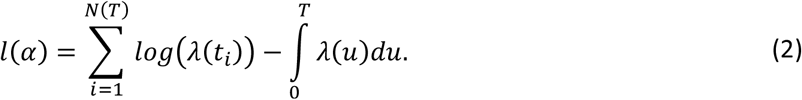

The maximum likelihood estimate 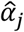 of *α*_*j*_ is obtained by solving

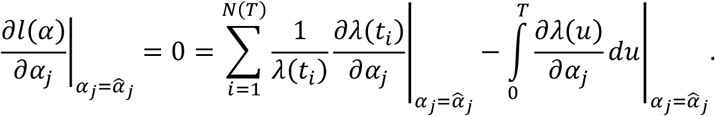

Using Eq. 1 we obtain

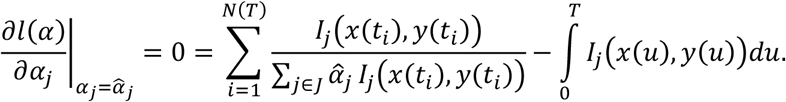

Let *s*_*j*_ denote the number of spikes that have occurred inside the *j*^th^ bin throughout the experiment. Since *I*_*j*_(*x*(*t*_*i*_), *y*(*t*_*i*_)) is non-zero if and only if (*x*(*t*_*i*_), *y*(*t*_*i*_)) is inside the *j*^th^ bin, the summation term simply gives 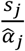. On the other hand, the integrand is non-zero if and only if (*x*(*u*), *y*(*u*)) is inside the *j*^th^ bin. Therefore, the integral gives the total amount of time spent in the *j*^th^ bin. Let that duration be denoted by *T*_*j*_. Then, the maximum likelihood estimate of *α*_*j*_ is obtained as

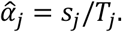

This is the ratio that Muller et al. computed to construct their firing rate maps (Muller et al., 1987). This shows that their firing rate maps are maximum likelihood estimates of the parameters of the conditional intensity function in Eq. 1.

Note that 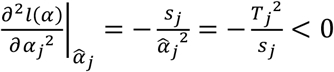 if *s*_*j*_ > 0, since *T*_*j*_ > 0 for visited bins, demonstrating that the estimate is indeed a maximum. On the other hand, if *s*_*j*_ = 0, then 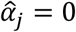 and since 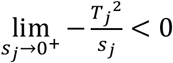, the log-likelihood has a maximum at 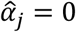.

#### 3.1.2 Model Selection

Using the maximized log-likelihood, Akaike Information Criterion can be computed for a firing rate map that has *K* visited bins, where *K* is the cardinality of *J* in Eq. 1. The Akaike Information Criterion that is corrected for sample sizes that are comparable to the number of parameters is (Burnham & Anderson, 2002; Sugiura, 1978)

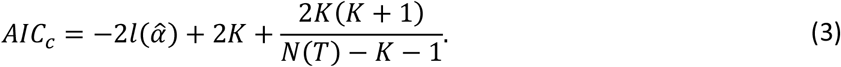

In Eq. 3, 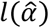 is computed using Eq. 2 as shown in Eq. 4:

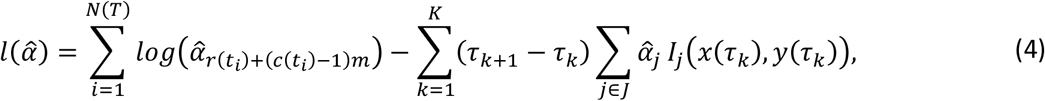

where, *τ*_*K*+1_ is taken as *T*. Since *I*_*j*_(*x*(*τ*_*k*_), *y*(*τ*_*k*_)) = 1 if the position sample (*x*(*τ*_*k*_), *y*(*τ*_*k*_)) is in the *j*^th^ bin and 0 otherwise, the double summation in Eq. 4 becomes 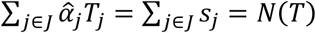, yielding

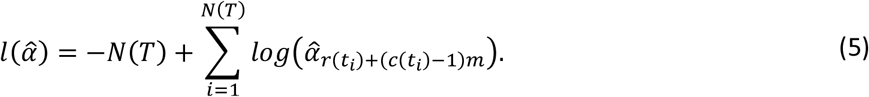

##### 3.1.2.1 Optimization of *m* and *n*

The spatial resolution of the firing rate map can be optimized by optimizing the numbers *m* and *n*. Let *λ*(*t*; *m, n*) denote the conditional intensity function model constructed using an *m*-by-*n* array of bins:

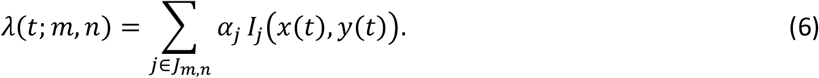

Let the *AIC*_*c*_ of this model be denoted by:

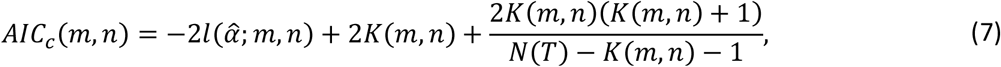

where *K*(*m, n*) is the number of visited bins at this spatial resolution.

*K*(*m, n*) is expected to increase with *m* and *n*, but the denominator *N*(*T*) − *K*(*m, n*) − 1 enforces the inequality

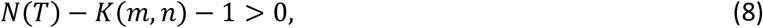

which limits the maximum numbers that can be used for *m* or *n*. Then the optimal values of *m* and *n* are

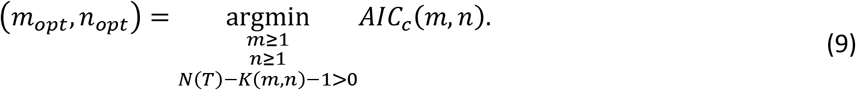

##### 3.1.2.2 Model Averaging

Let Δ*AIC*_*c*_(*m, n*) denote the difference *AIC*_*c*_(*m, n*) − *AIC*_*c*_(*m*_*opt*_, *n*_*opt*_) for a given cell. For some cells, Δ*AIC*_*c*_(*m, n*) ≤ 4 may hold for more than one pair of (*m, n*) values. Models with Δ*AIC*_*c*_ ≤ 4 are considered to be comparable in performance to the *AIC*_*c*_-best model (Burnham & Anderson, 2002; Cavanaugh & Neath, 2019; Portet, 2020). In such cases, the conditional intensity function of the place cell may be better represented by Akaike-weighted average of the comparable firing rate maps. Let *q* index the (*m*_*q*_, *n*_*q*_) pairs that belong to models for which Δ*AIC*_*c*_(*m*_*q*_, *n*_*q*_) ≤ 4, *q=*1, 2, …, Q. Then, Akaike-weighted average firing rate map is defined as

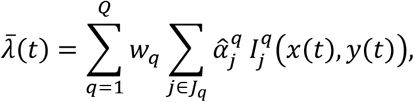

where 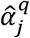 is the firing rate estimate in bin *j* of map *q*, 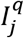 is the indicator function for that bin, and the Akaike weight *w*_*q*_ of model *q* is defined as (Burnham & Anderson, 2002, 2004; Portet, 2020)

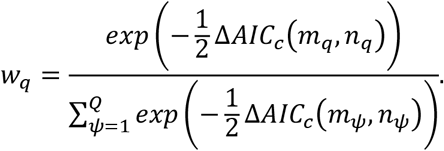

Since the firing rate maps are *m* by *n* matrices, Akaike-weighted average map is constructed as a *μ* by *υ* matrix, where *μ* is the least common multiple of *m*_*q*_, and *υ* is the least common multiple of *n*_*q*_, *q=*1, 2, …, Q.

#### 3.1.3 Absolute Goodness-of-Fit Test

According to the Time-Rescaling Theorem, the true conditional intensity function that underlies the spike train can be used to rescale the inter-spike intervals (ISIs), converting the spike train into a homogeneous Poisson process with unit rate (Poisson 1) (Brown et al., 2002; Ogata, 1988; Papangelou, 1972). The latter has the properties that its ISIs are mutually independent and exponentially distributed with unit mean. When the optimal firing rate map is used in place of the true conditional intensity function, the closeness of the resulting re-scaled spike train to Poisson 1 serves as a measure of how well the firing rate map approximates the true conditional intensity function underlying the observed spike train.

To compare the resulting re-scaled spike train to Poisson 1, first, the Probability Integral Transform is used for transforming the re-scaled ISIs into a random variable, *u*, that would have a standard Uniform distribution if the firing rate map were identical to the true conditional intensity function. The distribution of *u* is compared against the standard Uniform distribution using the Kolmogorov-Smirnov (KS) test (Brown et al., 2002). Next, the Inverse Probability Integral Transform is used to transform *u* into a new random variable, *z* = Φ^−1^(*u*), that would have a standard Normal distribution if the firing rate map were identical to the true conditional intensity function. Here, Φ^−1^(·) is the inverse of the standard Normal cumulative distribution function. The autocorrelation of these re-scaled and double-transformed ISIs (*z* values) is computed at all possible lags. If the result of autocorrelation is consistent with zero-mean white noise with variance *φ*^−1^, where *φ* is the number of autocorrelated *z* values, then it is concluded that the re-scaled ISIs are mutually independent (Czanner et al., 2008). This is based on the fact that uncorrelatedness implies independence for Normal r.v.s. (Box et al., 2016, p. 33).

In the present study, the analyses were performed in MATLAB (MathWorks, Natick, MA, USA). In computing the autocorrelation, the value of the random variable *z* may become infinity (Inf in MATLAB) for some samples. Since *z* = Φ^−1^(*u*), such Inf values are obtained because *u* is 1 for those samples. The latter is due in turn to the presence of re-scaled interspike intervals that are too long for a unit Poisson process, indicating that the firing rate map fails to successfully re-scale the corresponding interspike intervals. Theoretically, since *u* = 1 can only be obtained for an interval that is infinitely long, the occurrence of this value during an analysis is due to finite computing precision. To avoid it, the value of such *z* samples were set to 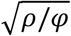, where *ρ* = 1.7977 × 10^308^ is the largest positive floating-point number in MATLAB. In this way, the autocorrelation was ensured to never exceed the finite value of *ρ* even when every single *z* sample that enters the autocorrelation equals 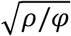.

### 3.2 Features Extracted from Firing Rate Maps

Previous studies extracted various features from firing rate maps to quantify the characteristics of how place cell firing relates to behavior and cognition. Some of those features include the number, shape, area (the number of bins that make up a place field, i.e. the region in which the cell fires) and location (location of peak firing rate) of the place fields within the experimental apparatus (Hussaini et al., 2011; Muller et al., 1987), patchiness (the number of distinct bin groups in which the firing rate falls within a fixed range) (Muller & Kubie, 1989), coherence (the z-transform of the Pearson’s correlation coefficient R between a list of firing rates in each bin and a corresponding list of firing rates averaged over the 8 nearest neighbors of each bin, where the z-transform refers to Z = 0.5 ln ((1 + R)/(1 − R)) (Bures et al., 1997; Hussaini et al., 2011; Muller & Kubie, 1989; Zhang et al., 2014), concentration (the ratio of the number of bins in the place field to the total number of bins where a spike was detected), dispersion (the number of bins in which a spike was detected divided by the apparatus area) (Bures et al., 1997), and information content (Hussaini et al., 2011; Skaggs et al., 1992) of the place field.

In identifying the place fields, a firing rate map is first thresholded at a level determined by the experimenter, such as 1 Hz (Muller et al., 1987). Next, groups of contiguous pixels are identified. The contiguity condition is usually the sharing of at least one edge with another place field bin (Muller et al., 1987). Thus, starting from the suprathreshold local maxima of the firing rate map, place fields can be constructed iteratively until no bin can be adjoined to existing place fields. Usually, a threshold can also be defined for the minimum place field size, such as 9 bins (Muller et al., 1987).

These methods may be adapted to accommodate the optimal firing rate maps obtained using the point process framework presented here. Comments are provided in the Discussion as to how these features can be computed in the point process framework and how, in some cases, these features may be replaced by other tools that this framework offers to reach the same end.

### 3.4 Comparison of Results Between Rats

After *m*_*opt*_ and *n*_*opt*_ values are determined for all cells of the two rats, the non-parametric Kruskal-Wallis (KW) test is used for testing whether they are significantly different within or between rats, supplemented by Tukey-Kramer multiple comparison test. The KW test is also used for testing whether *K*(*m*_*opt*_, *n*_*opt*_) values differ between the two rats.

In addition, KS-test P values are compared between the two rats using the Wilcoxon rank sum test. Also, the relation between the logarithm of the KS-test P value and the logarithm of the session-average firing rate is explored using multiple regression, where the former is fit as a function of a power series expansion in the latter.

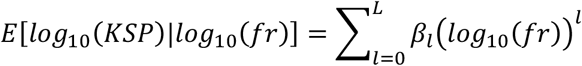

The optimal order *L* of the power series expansion is determined by minimizing the *AIC*_*c*_ of the multiple regression model. Model averaging is performed using Akaike weights if two or more comparable regression fits are obtained.

## 4. Results

In this section the *AIC*_*c*_(*m, n*) plot is illustrated for a representetative cell. Summary statistics are reported for the optimal (*m, n*) values of all cells for both rats. Plots of optimal and model-averaged firing rate maps are provided for representative cells. Absolute goodness-of-fit test results are provided. Results are compared within and between rats.

### 4.1 Optimization of *m* and *n*

To speed up the optimization of *m and n, AIC*_*c*_(*m, n*) is first minimized under the constraint *m* = *n* = *u*. Let *u*_*opt*_ denote the value that minimizes *AIC*_*c*_(*u, u*) (e.g. Fig. 1A). Then, the optimal values of *m* and *n* are searched within the region *max*(1, *u*_*opt*_ − 20) ≤ *m* ≤ *u*_*opt*_ + 20 and *max*(1, *u*_*opt*_ − 20) ≤ *n* ≤ *u*_*opt*_ + 20, subject to the inequality 8. Except for cell 7 of rat 1, the pair (*m*_*opt*_, *n*_*opt*_) was found within this search space, never being located on the upper boundaries (e.g. Fig. 1B). For cell 7 of rat 1, the search space was expanded to *max*(1, *u*_*opt*_ − 30) ≤ *m* ≤ *u*_*opt*_ + 30 and *max*(1, *u*_*opt*_ − 30) ≤ *n* ≤ *u*_*opt*_ + 30, and (*m*_*opt*_, *n*_*opt*_) was found within this search space, without being located on the upper boundaries.

**Figure 1.**
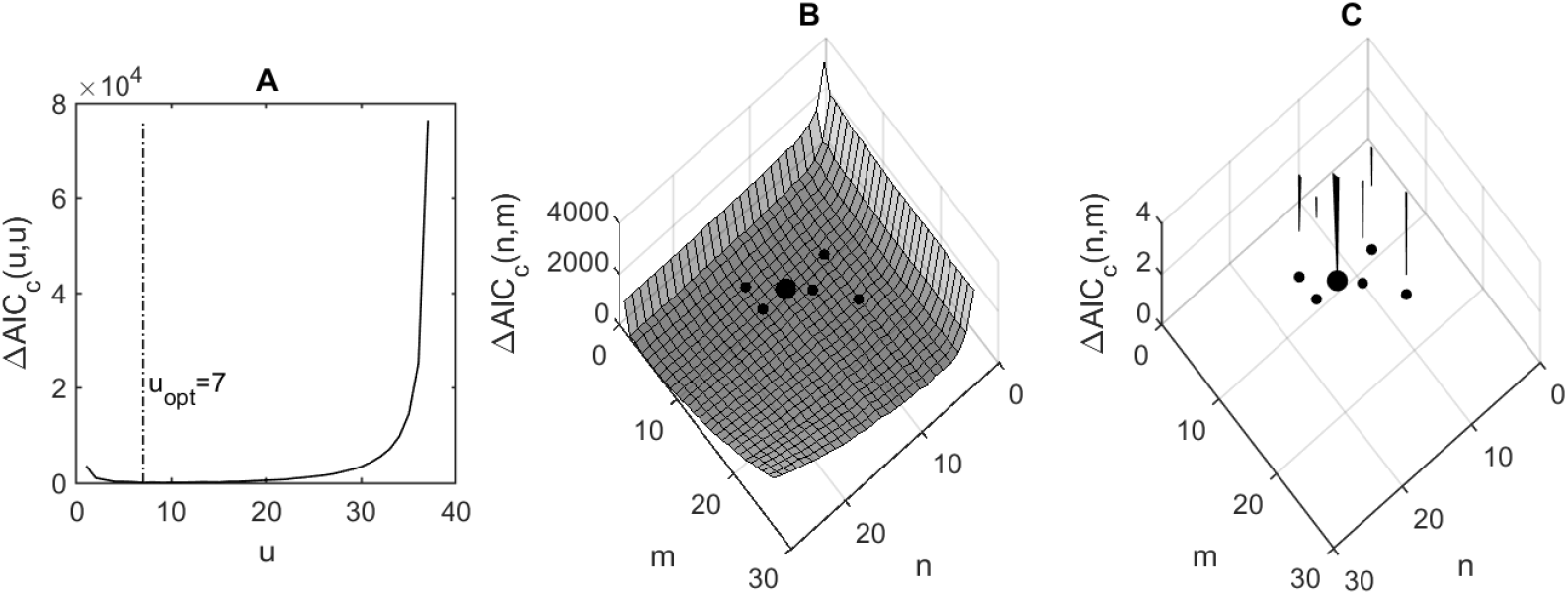
Optimization of the size and the aspect ratio of histogram bins. (A) *AIC*_*c*_(*u, u*) was minimized at *u*_*opt*_ = 7 for cell 1 of rat 1. (B) *AIC*_*c*_-best firing rate map was found at (*m*_*opt*_, *n*_*opt*_) = (9,12) (large disk). Δ*AIC*_*c*_(*m, n*) ≤ 4 was satisfied at five other (*m, n*) values (small disks). (C) The part of the Δ*AIC*_*c*_(*m, n*) surface that lies below the level Δ*AIC*_*c*_(*m, n*) = 4 is shown.

The optimal values of *m* and *n* ranged between 1 and 33 across all cells considered (Table I). The optimal map obtained in Fig. 1 is illustrated in Fig. 2. KW test did not detect any significant difference between *m*_*opt*_ and *n*_*opt*_ values within or between the two rats (H(3)=5.34, P=0.14). However, there was a significant difference in the total number of bins per firing rate map (*m*_*opt*_ × *n*_*opt*_) between the two rats (H(1)=5.03, P=0.025). Similarly, *K*(*m*_*opt*_, *n*_*opt*_) values were also different (H(1)=4.93, P=0.0264). Consistent with these findings, KW test did not detect any significant difference between the optimal bin width X versus Y values within or between the two rats (H(3)=4.89, P=0.18), yet there was a significant difference in the optimal bin area (*XY*) between the two rats (H(1)=4.15, P=0.0417).

**Table I.**
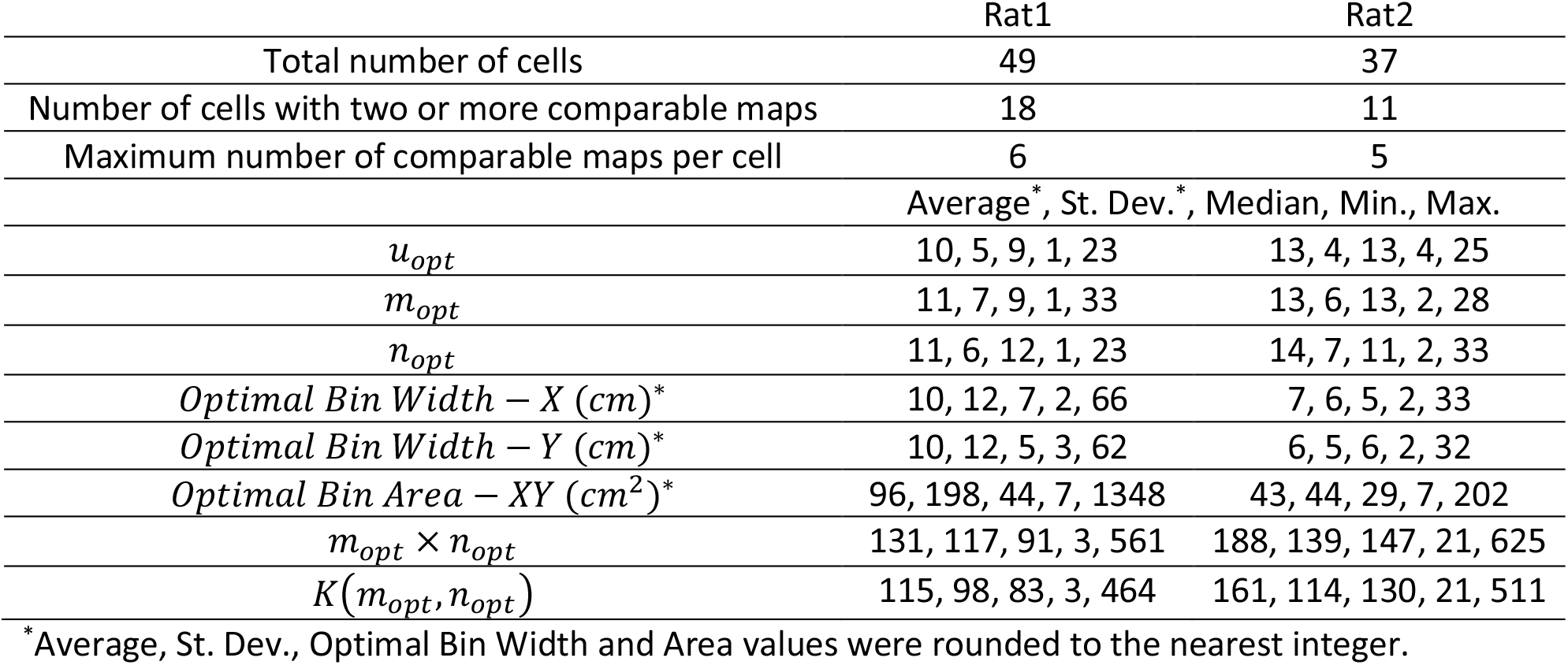
Optimal bin size summary.

|  | Rat1 | Rat2 |
| --- | --- | --- |
| Total number of cells | 49 | 37 |
| Number of cells with two or more comparable maps | 18 | 11 |
| Maximum number of comparable maps per cell | 6 | 5 |
|  | Average*, St. Dev.*, Median, Min., Max. |  |
| $u_{opt}$ | 10, 5, 9, 1, 23 | 13, 4, 13, 4, 25 |
| $m_{opt}$ | 11, 7, 9, 1, 33 | 13, 6, 13, 2, 28 |
| $n_{opt}$ | 11, 6, 12, 1, 23 | 14, 7, 11, 2, 33 |
| Optimal Bin Width – X (cm)* | 10, 12, 7, 2, 66 | 7, 6, 5, 2, 33 |
| Optimal Bin Width – Y (cm)* | 10, 12, 5, 3, 62 | 6, 5, 6, 2, 32 |
| Optimal Bin Area – XY (cm<sup>2</sup>)* | 96, 198, 44, 7, 1348 | 43, 44, 29, 7, 202 |
| $m_{opt} \times n_{opt}$ | 131, 117, 91, 3, 561 | 188, 139, 147, 21, 625 |
| $K(m_{opt}, n_{opt})$ | 115, 98, 83, 3, 464 | 161, 114, 130, 21, 511 |

\*Average, St. Dev., Optimal Bin Width and Area values were rounded to the nearest integer.

**Figure 2.**
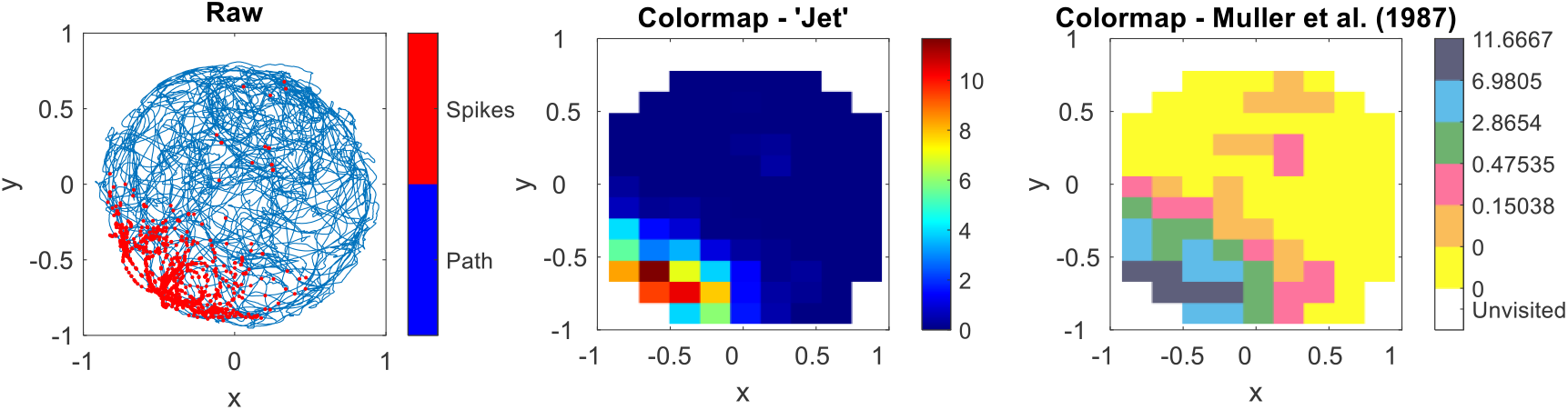
Optimal firing rate map. The raw data show the spike locations of cell 1 of rat 1 as red dots on the trajectory of the rat’s head (blue trace). The optimal firing rate map obtained in Fig. 1 is shown using two different coloring schemes. MATLAB’s colormap called ‘Jet’ reveals the firing rate gradient with continuous color levels. The coloring scheme of Muller et al. quantizes the firing rate as shown in the colorbar (Muller et al., 1987). The unvisited bins are shown in white, 0 Hz is shown in yellow, maximum firing rates in the other colors are indicated on the upper edge of each color segment.

These differences indicate that more parameters per unit area were required to represent the position dependence of rat 2 place cell firing. This suggests that more position information was encoded in rat 2 place cell spike trains. To check whether this was associated with any difference in the firing rates of the place cells of the two animals, the session-average firing rates of the place cells were compared. The median value of the session-average firing rate of the rat 2 place cells was twice that of rat 1’s (1.26 Hz versus 0.6 Hz; P=0.0129; Wicoxon ranksum test).

### 4.2 Model Averaging

Two or more pairs of (*m, n*) values provided comparable fits to the *AIC*_*c*_-best firing rate map in 18 and 11 cells in rats 1 and 2, respectively (e.g. Fig. 1C). The relative numbers of these cells were not significantly different between the two rats (P>0.64, Fisher’s exact test). For such cells, Akaike-weighted average firing rate map was computed (e.g. Fig. 3).

**Figure 3.**
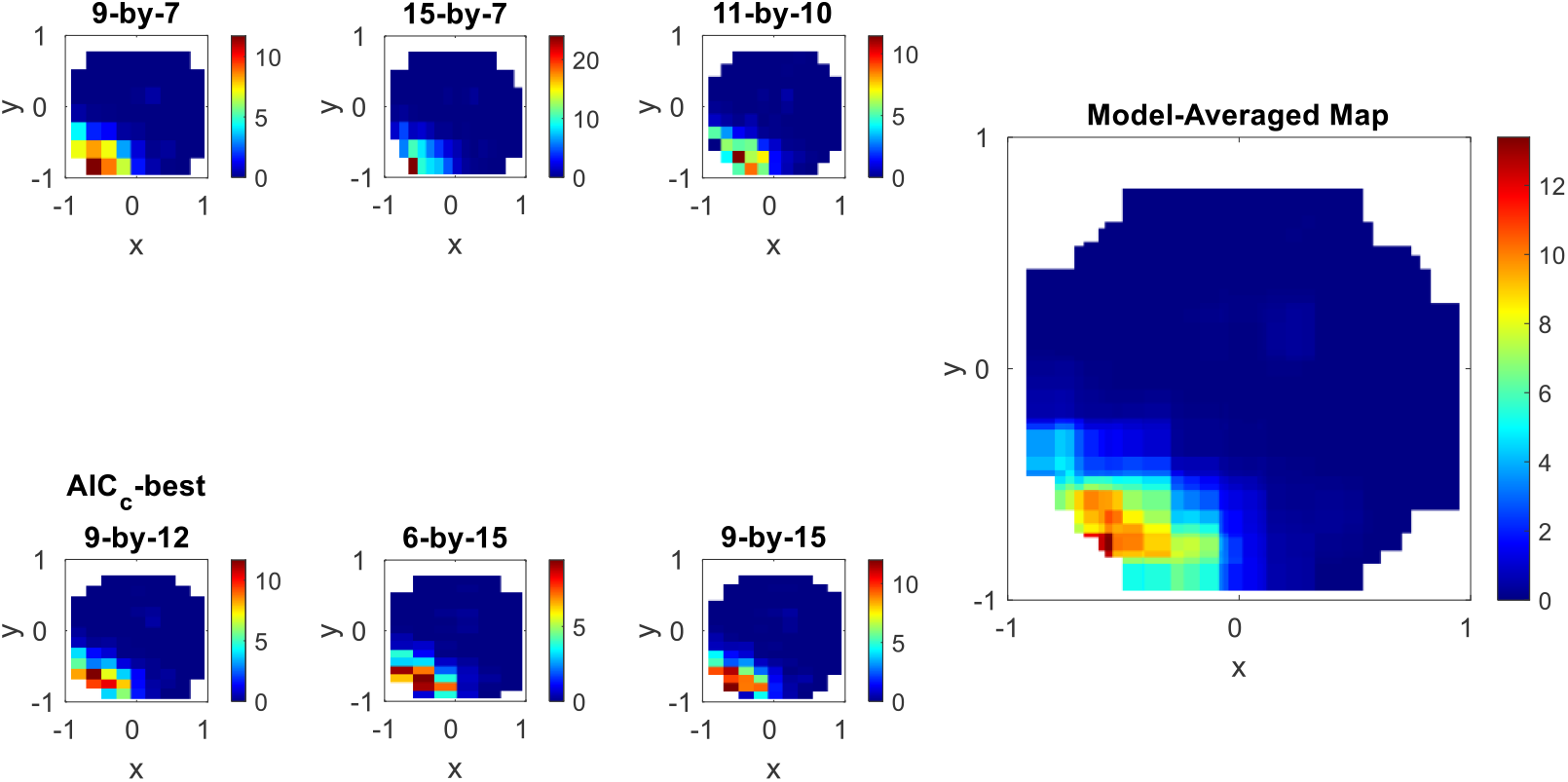
Model averaging using Akaike weights. The six smaller panels on the left show the firing rate maps that have Δ*AIC*_*c*_(*m, n*) ≤ 4 in Fig. 1. Since they are comparable in relative goodness-of-fit, the firing rate map may be better estimated by taking all of them into account using Akaike-weights. The model-averaged map is shown on the right as the larger panel. The KS test P-value of the *AIC*_*c*_-best map and the model-averaged map were 5.96 10^−29^ and 3.87 10^−27^, respectively, indicating that model averaging improved the fit 65-fold.

### 4.3 Absolute Goodness-of-Fit Test

Only the optimal firing rate map of cell 38 (rat 1) passed the absolute goodness-of-fit test (Fig. 4A and 4D). KS-test P-values ranged between 10^−282^ and 10^−5.6^ for the other cells (Fig. 4) and were significantly smaller for rat 2 (median values: 10^−113^ vs 10^−55^, P=0.01, Wilcoxon rank sum test). The optimal firing rate maps of the cells in Fig. 4 are shown in Fig. 5.

**Figure 4.**
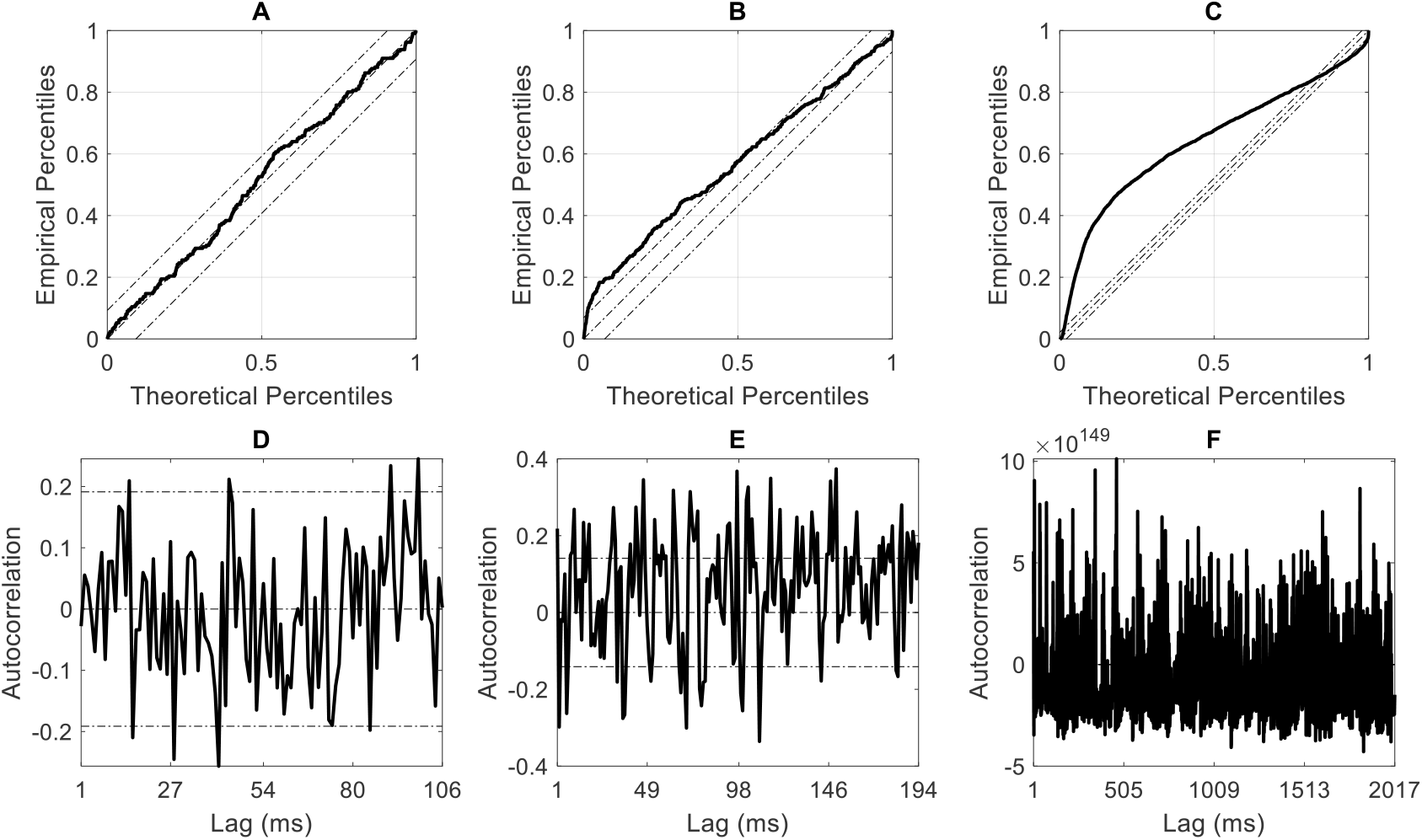
The best and the worst KS plots and associated autocorrelation plots. (A and D) The optimal firing rate map of cell 38 of rat 1 is the only optimal model that passed the absolute goodness-of-fit test (KS-test P=0.38). (B and E) The next best test result was obtained for cell 20 of rat 1 (KS-test P=10^−5.58^). (C and F) The optimal firing rate map that had the worst test result was obtained for cell 28 of rat 2 (KS-test P=10^−282^).

**Figure 5.**
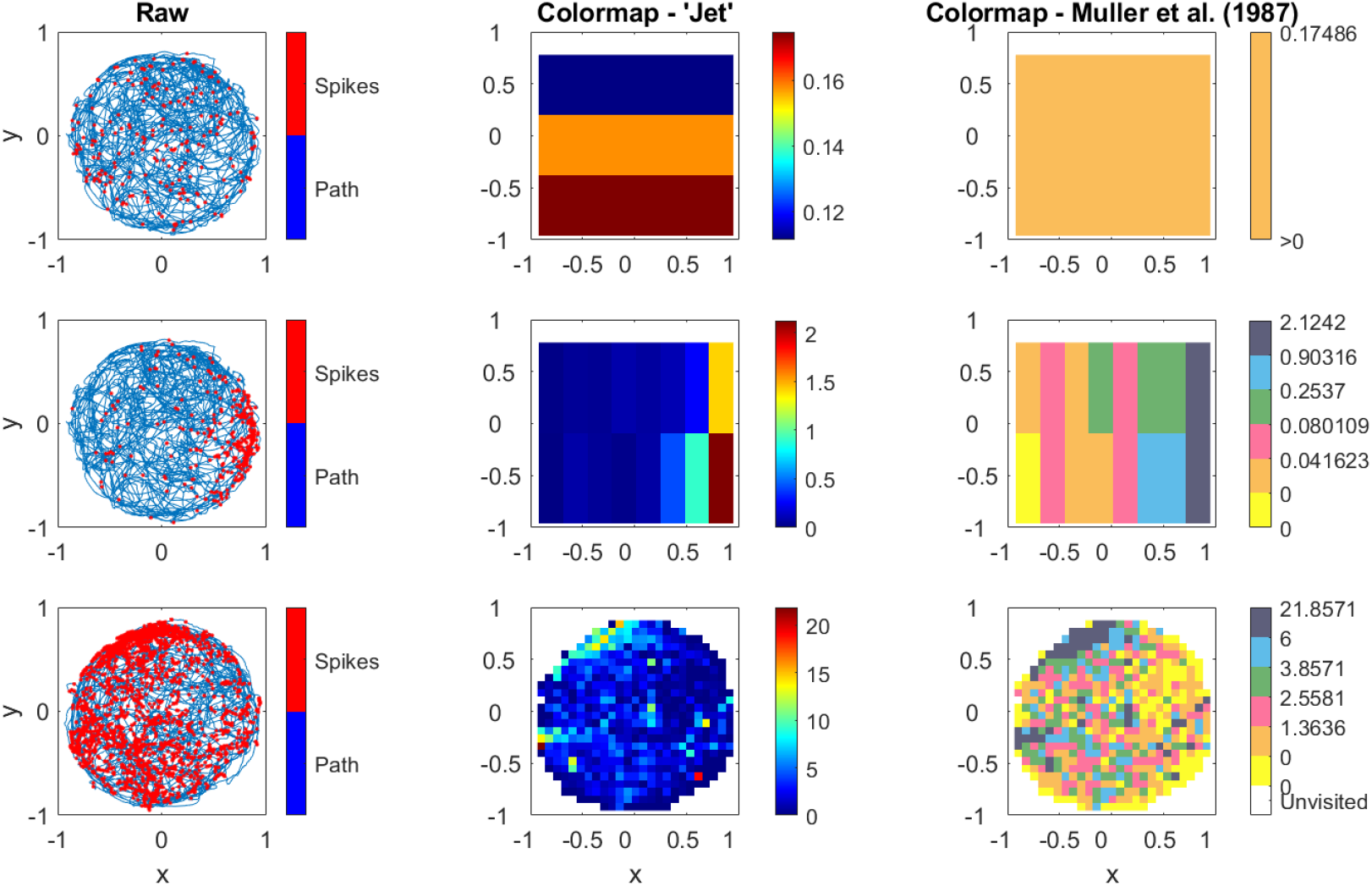
The optimal firing rate maps of cells in Fig. 4. Maps for cells 38, 20 and 28 are shown in the top, middle and bottom rows, respectively. Cell 38 (rat 1) (top) had a very low firing rate and fired virtually homogeneously throughout the environment. The optimal map had only 1 bin on the x-axis and 3 bins on the y-axis. The coloring scheme of Muller et al. (1987) represents this activity using only one color (From Table 2 Map E of ref.). Cell 20 (rat 1) (middle) seemed to have a weak dependence on position in the vertical orientation. As a result, the optimal map had only two bins on the y-axis. Because each bin of the firing rate map was visited by the rat, the map reflects the rectangular structure of the firig rate map as a 2-D matrix and covers the entire trajectory for cells 38 (top) and 20 (middle). Cell 28 (rat 2) (bottom) reached firing rates of up to about 22 Hz in some bins. The optimal firing rate map had 25 bins on both the x- and y-axes.

Because the session-average firing rate was found to be significantly higher for rat 2 (see Section 4.1), it was hypothesized that the difference between the KS-test P-values between the two rats might be related to the difference between the session-average firing rates. Supporting this hypothesis, multiple regression analysis (see Section 3.4) revealed that KS-test P-values were progressively lower as the session-average firing rate increased (Fig. 6).

**Figure 6.**
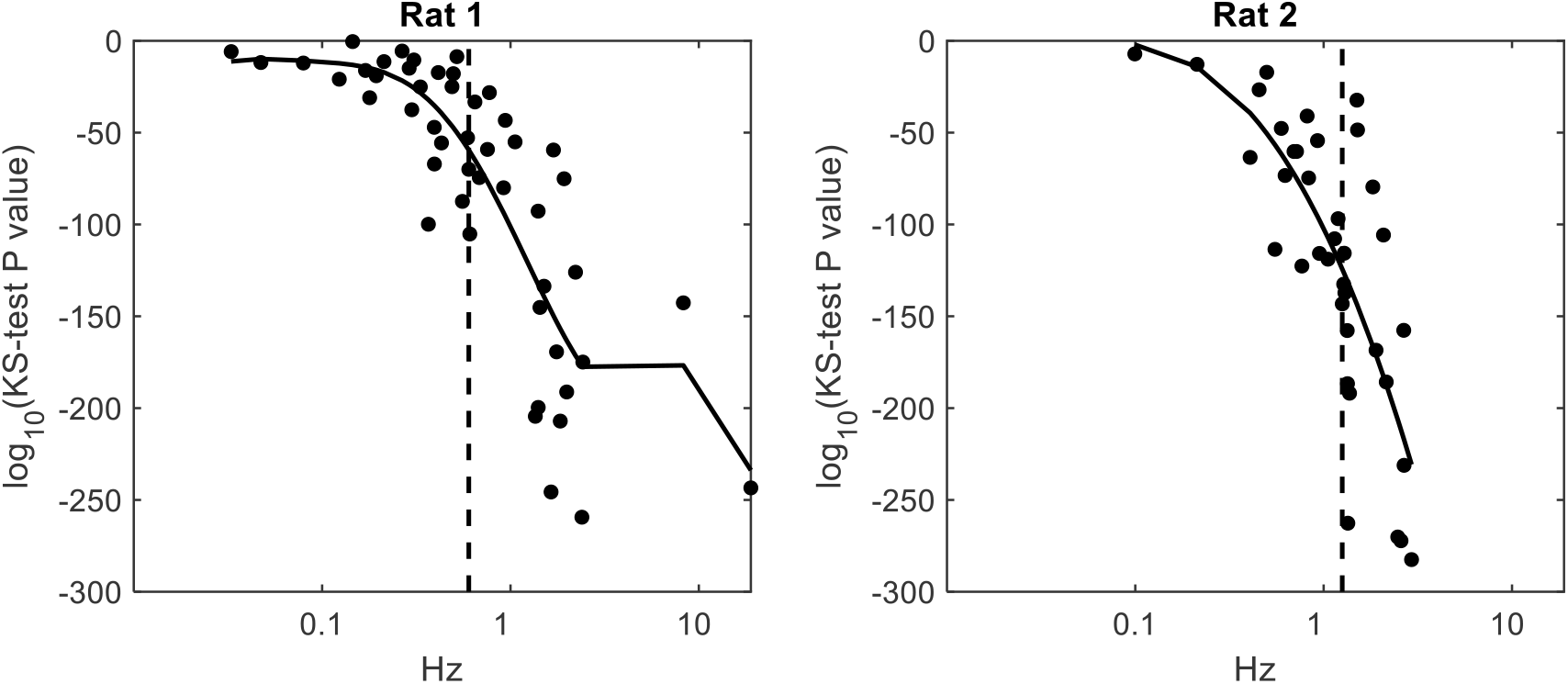
Firing rate map absolute goodness-of-fit deteriorates with increasing session-average firing rate. For rat 1, base-10 logarithm of the KS-test P-value was optimally fit by a 6^th^ order power series expansion in the base-10 logaritm of the session-average firing rate. Model orders 4 through 7 provided comparable fits. In all those models the first order slope was negative and significant (P<2 10^−7^). For rat 2, the optimal fit was provided by a second order power series expansion, and model orders 1 and 3 provided comparable fits. In all those models the first order slope was negative and significant (P<2 10^− 4^). Model-averaged fits are shown for eact rat. Vertical dashed lines indicate the median value of the session-average firing rate.

#### 4.3.1 Absolute Goodness-of-Fit of Model-Averaged Maps versus AIC_c_-best Maps

Of the 29 cells for which two or more maps provided comparable fits to the data, the model-averaged firing rate map achieved a higher (better) KS-test P-value than the *AIC*_*c*_-best model in 17 cases. This improvement was observed in more cells of rat 1 (14/18) than rat 2 (3/11) (P=0.018; Fisher’s exact test), suggesting that the improvement occurred more frequently at low session-average firing rates. Indeed, the mean session-average firing rate was 0.64 Hz (0.85 Hz) in the 14 (3) cases where model averaging provided better fits, compared to 4.87 Hz (1.15 Hz) in the remaining 4 (11) cases for rat 1 (rat 2). However, these within rat differences did not reach statistical significance (P>0.08; Two-sample t-test), possibly due to the small sample sizes. Six different firing rate maps were found to be comparable for cell 38 of rat 1. The model-averaged map passed the absolute goodness-of-fit test for that cell (P>0.47; KS-test). In all other cases, the model-averaged maps failed the absolute goodness-of-fit test (P<10^−5.28^, KS-test).

## 5. Discussion

Firing rate maps that are constructed using 2-D spike-in-location and time-in-location histograms have been one of the earliest, simplest and most widely used methods for quantifying the position dependence of place cell spiking (Ásgeirsdóttir et al., 2020; Bures et al., 1997; Fenton & Muller, 1996, 1998; Harvey et al., 2020; Hazama & Tamura, 2019; Hussaini et al., 2011; Jung et al., 1994; Karnam et al., 2009; Muller, 1996; Muller et al., 1987, 1994; O’Keefe, 1983; Sakkaki et al., 2020; Schoenenberger et al., 2016; Skaggs & McNaughton, 1998; Solstad et al., 2014; Tanila et al., 2018; Zhou et al., 2007). Few attempts have been made to optimize the firing rate estimates computed using these maps, and these attempts neither optimized the number of bins used per cell, nor did they use the point process framework (Grieves, 2023; Jung et al., 1994; Skaggs et al., 1996; Yartsev & Ulanovsky, 2013).

One of the methods that aimed to optimize the firing rate map is the “adaptive binning” method (Jung et al., 1994; Skaggs et al., 1996). In this method the experimental area was divided into a fixed number of bins (64×64), and the firing rate in bin *j* was estimated as the ratio of the total number of spikes observed in a circular area of radius *r*_*j*_, centered at bin *j*, divided by the total duration spent in that circular area. The radius *r*_*j*_ was the smallest radius that was greater than or equal to 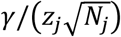, where *γ* was called the “sampling radius scale factor” and was set to 1000, *z*_*j*_ was the total time spent in the circular area, and *N*_*j*_ was the total number of spikes observed in that area. As a result, the firing rate maps had the same number of bins for each place cell, and the firing rate in each bin was estimated using data from a circular area with a radius that varied from bin to bin. This approach does not use the point process framework since the firing rate it computes was not optimized by maximizing the point process likelihood function.

In a related method, the radius *r*_*j*_ was determined as the radius of the smallest circular area, centered at bin *j*, in which the total time-in-location, *z*_*j*_, was at least 1 s (Yartsev & Ulanovsky, 2013). The firing rate estimate at bin *j* was then calculated as the ratio of the total number of spikes counted in that circular area, divided by *z*_*j*_. Clearly, this method too divided the space into the same number of bins for each place cell and did not use the point process framework to optimize the firing rates.

A recent simulation study tested the accuracy with which these firing rate map optimization methods estimate the true firing probability underlying the data (Grieves, 2023). Firing rate maps with square bins of various sizes were constructed using the adaptive binning methods described above. The mean integrated square error (MISE) between the estimated and the actual firing probability was used to quantify the rate map error. The smallest MISE values were obtained for the smallest bin size possible (<0.5 cm) and a relatively large amount of smoothing (large *r*_*j*_). Optimal bin size increased with the size of the simulated place field and decreased with recording duration. Low data densities were found to benefit from larger bins, as was also observed in the present study (e.g. rat 1 vs rat 2 in Table I). However, this study too did not use the point process framework to optimize the firing rate maps.

In contrast with these studies, in the present method, the point process log-likelihood function is constructed (Eq. 2), which provides the basis for optimizing all model parameters through *AIC*_*c*_-based model selection (Eq. 9). The absolute goodness-of-fit is assessed based on statistical tests developed using the Time-Rescaling Theorem (Sections 3.1.3 and 4.3). Moreover, models that provide fits comparable to the *AIC*_*c*_-best model are averaged using Akaike weights. Whether model averaging improves the absolute goodness-of-fit is assessed. Indeed, it was found that model averaging provided better fits for cells that had a relatively low session-average firing rate.

For a substantial number of cells, the optimal spatial resolution of the firing rate map was found to be much lower than what is usually used in the literature (Fig. 2, Fig. 3, Fig. 5, Table I). For half of the cells, the bin width (or height) was found to be more than 5 cm, which is larger than the bin widths of 4 cm (Schoenenberger et al., 2016), 3 cm (Fenton & Muller, 1996, 1998; Harvey et al., 2020; Hayman et al., 2015; Ismakov et al., 2017), 2.3-2.7 cm (Jung et al., 1994; Karnam et al., 2009; Muller, 1996; Muller et al., 1994; Zhou et al., 2007), 2 cm (Sakkaki et al., 2020; Skaggs & McNaughton, 1998; Wikenheiser et al., 2021), 0.9-1 cm (Ásgeirsdóttir et al., 2020; Hazama & Tamura, 2019), and 0.5 cm (Grieves, 2023) that are encountered in the literature. This suggests that studies that use higher spatial resolutions in histogram-based firing rate maps may be overfitting the data for some cells.

The new firing rate map construction method presented here may necessitate revising the existing methods for extracting features from firing rate maps and place fields. Most conspicuously, in the proposed method, the “size” or “area” of a place field needs to be measured not in the number of bins it contains but in cm squares or as a fraction of the total area of the apparatus. That’s because the bin size may differ from one cell’s firing rate map to another’s in the present method. The same adjustment also applies to the computation of “concentration” and “dispersion”. Similarly, in the proposed method, the “location” of a place field can no longer be determined by identifying the bin with the highest firing rate. Rather, the coordinates of the center of the bin with the highest firing rate, relative to an origin fixed on the apparatus, would have to be computed. Whilst existing methods for the determination of “patchiness” or “information content” may continue to apply to the optimal firing rates produced by the point process framework, this may not be so for the computation of “coherence” when the firing rate map consists only of a few bins. A solution may be to sample firing rate values on a fixed spatial grid placed on the optimal firing rate maps and to compute the coherence using those samples.

Place field features are often used for addressing questions about the stability of place fields in different experimental conditions (Hussaini et al., 2011), or for estimating additional properties of position encoding, such as the optimal latency between the spike trains and the position time series (Muller & Kubie, 1989). Place field stability is quantified using the correlation coefficient between different firing rate maps. With maps that contain different numbers of bins, the correlation coefficient can be computed using the approach proposed for the computation of coherence. Namely, firing rate values may be sampled on a fixed spatial grid placed on the optimal firing rate maps, and the correlation coefficient may be computed using those samples. For the estimation of the optimal latency between the spike trains and the position time series, area, patchiness and coherence were optimized as a function of the latency (Muller & Kubie, 1989). By contrast, within the point process framework, the same estimation can be performed using the profile log-likelihood function of the latency (Okatan, Frank, et al., 2005). Therefore, the use of three features can be replaced by the use of one likelihood-based estimation method, which also provides a 95% confidence interval for the latency of individual or populations of cells (Okatan, Frank, et al., 2005). Overall, existing methods of extracting features from the firing rate maps can be used with the optimal maps produced by the present method, either as they are or with minor adaptations, while the point process framework may offer new approaches that can replace the existing methods in certain cases.

In some studies, firing rate maps are smoothed by convolving the spike-in-location and the time-in-location maps with a Gaussian kernel before taking their ratio (Grieves, 2023). In some other studies, smoothing is performed on the firing rate map, after the ratio of the spike map to the time-in-location map is computed (Harvey et al., 2020; Wang et al., 2020; Zutshi et al., 2018). Such methods have the drawback of artificially propagating spike-in-location, time-in-location or firing rate values to unvisited pixels, which then needs to be corrected by taking further measures (Grieves, 2023). Another issue concerns the selection of the scale of the smoothing kernel, which is often subjective. In the present method, no explicit smoothing is performed on the spike-in-location, time-in-location or the firing rate maps. Instead, if multiple firing rate maps of various bin sizes are found to provide comparable fits to the data, then they can be averaged using Akaike weights, which provides smoother firing rate maps (Fig. 3). We found, however, that such averaging improves the goodness-of-fit only in cases where the session-average firing rate is relatively low.

A further method that is used for constructing smooth firing rate maps is the average shifted histogram (ASH) method. In that method multiple firing rate maps are constructed with the same bin size by shifting the bin edges by fractions of the bin width in one or both directions; then, these multiple firing rate maps are averaged to produce a smoother map (Scott, 1985, 2015; Scott & Thompson, 1983). Usually the maps are averaged with equal weights (Bourel et al., 2014; Scott, 1985; Scott & Thompson, 1983), though non-equal weigths may also be used subject to certain constraints (Scott, 1985). By contrast, smoothing is performed in the point process framework through model averaging using Akaike weights, provided that multiple maps are found to fit the data comparably to the *AIC*_*c*_-best map. Thus, smoothing is performed as an entirely data driven procedure that does not make use of any free parameters. The point process framework could also accommodate the ASH method by computing the *AIC*_*c*_ of the shifted maps and including into model averaging those that provide fits comparable to the *AIC*_*c*_-best model. However, this was not implemented in the present analysis.

In the present analysis, the optimal firing rate map passed the absolute goodness-of-fit test for only one cell (cell 38 of rat 1, Fig. 4 and Fig. 5). Absolute goodness-of-fit deteriorated with increasing session-average firing rate (Fig. 6). This is not surprising in view of the fact that the spike trains of place cells are known to contain other information, in addition to position, such as the phase of the hippocampal theta rhythm (Skaggs et al., 1996), head direction, running speed, (McNaughton et al., 1983) acceleration (Kropff et al., 2021), functional connectivity (Okatan, Wilson, et al., 2005), and reward (Hölscher et al., 2003). These additional variables may need to be properly represented within the conditional intensity function models for the absolute goodness-of-fit test to be passed. Only the conditional intensity functions that pass the absolute goodness-of-fit tests can be trusted to provide an accurate representation of the position dependence of place cells.

Firing rate maps have also been constructed using other position dependent conditional intensity function models, such as the bivariate Gaussian (Brown et al., 1998) and expansions of polynomials, such as the Zernike polynomials (Barbieri, 2004; Barbieri et al., 2002) and bivariate power series (Margham & Okatan, 2023; Okatan et al., 2006). Compared to histogram-based firing rate maps, these approaches have the benefit of providing a conditional intensity function that is continuous in space. The optimal orders of the Zernike polynomial expansion and power series expansion models were previously determined for all cells of rat 2 (Margham & Okatan, 2023). For the Zernike polynomial expansion models of optimal order, the number of parameters used per cell ranged between 10 and 351, with a median value of 120, while these numbers were 9, 299 (incorrectly written as 340 in (Margham & Okatan, 2023)), and 135, respectively, for the power series expansion models of optimal order (Margham & Okatan, 2023). In comparison, the number of parameters used per cell ranged between 21 and 511, with a median value of 130, for the optimal firing rate maps (Table I). The detailed comparison of the relative and absolute goodness-of-fit of these different model types will be the subject of a future study.

The method presented here revealed that as simple as a 1-by-3 firing rate map suffices to model the firing rate of some place cells (Table I, Fig. 5-top). A higher spatial resolution would overfit such data according to *AIC*_*c*_. In the traditional approach, where a fixed number of bins is used for each cell, such a cell would have non-zero firing rate in some bins, and those bins would be distributed throughout the environment (e.g. as in Fig. 5E of (Muller et al., 1987)). The new representation of the firing rate makes it obvious that no detailed binning is necessary for such a cell. On the other hand, for cells that exhibit significant firing rate modulation with position, the new method discovers the optimal spatial resolution (e.g. Fig. 5-bottom) in the orientation in which it is needed (Fig. 5-middle).

For some cells, the orientation of the place field may not aling perfectly with the chosen vertical and horizontal axes of the environment. The firing field of the cell in Fig. 3, for instance, is located in the lower-left quadrant of the environment. The optimal numbers of bins were found to be 12 and 9 on the X and Y axes, respectively. If the data were rotated by about 45 degrees clockwise before constructing the firing rate map, then position dependence would be localized on the horizontal axis. In that case, the optimal firing rate might have had only a few bins on the vertical axis and more bins on the horizontal axis, as in the middle panel of Fig. 5. Thus, optimizing the orientation of the firing rate map with respect to the environment using a non-parametric method, such as the principle components analysis, may further improve *AIC*_*c*_. This will be tested in future studies.

The present method of firing rate map optimization revealed that more bins per unit area were needed to model the activity of the place cells of rat 2. This might not have been readily noticed in methods that use a fixed number of bins per firing rate map. However, this might have been detected in adaptive binning methods, where the values of *r*_*j*_ might have been smaller in rat 2. The reason why a higher spatial resolution was needed to model the place cells of rat 2 seems to be the higher session-average firing rates of those cells, compared to the cells of rat 1 (a ratio of about 2:1). If the firing rate of a place cell is a continuous non-uniform function of position, as most seem to be (e.g. Fig. 2), then, doubling that firing rate would indeed justify the use of more bins per unit area to represent the position dependent change in firing rate. Indeed, as was indicated earlier, simulation studies also show that low data densities benefit from larger bins (smaller number of bins per unit area) (Grieves, 2023).

Histogram-based firing rate maps are widely used for quantifying the position dependence of the place cell firing (Ásgeirsdóttir et al., 2020; Bures et al., 1997; Fenton & Muller, 1996, 1998; Grieves, 2023; Harvey et al., 2020; Hazama & Tamura, 2019; Hussaini et al., 2011; Jung et al., 1994; Karnam et al., 2009; Muller, 1996; Muller et al., 1987, 1994; Muller & Kubie, 1989; Sakkaki et al., 2020; Schoenenberger et al., 2016; Skaggs & McNaughton, 1998; Solstad et al., 2014; Tanila et al., 2018; Zhang et al., 2014; Zhou et al., 2007). The method presented here can be used to compute these maps by optimizing the size and the aspect ratio of the bins for individual place cells. The method also allows for the averaging of multiple maps using Akaike weights, and for the assessment of relative and absolute goodness-of-fit. Therefore, this method is expected to contribute to the study of the position dependent firing of hippocampal place cells. The proposed method can also be applied to the histogram-based quantification of the dependence of the firing rate on variables other than position.

## 6. Conclusions

Histogram-based firing rate maps of hippocampal place cells are optimized for each place cell using the point process framework. The optimal size and aspect ratio of histogram bins are obtained by minimizing the Akaike Information Criterion. Maps that have comparable relative goodness-of-fit are averaged using Akaike weights to obtain smoother maps. The absolute goodness-of-fit of the maps is assessed and found to be better for cells that have low session-average firing rate. Existing methods of feature extraction from firing rate maps are observed to also apply to the new optimized maps either directly or with minor adaptations. It is noted that the point process framework provides likelihood-based tools that are not available in existing methods of constructing histogram-based firing rate maps.

## Data availability statement

Data used in this study are available at Cajigas (2017).

